# A distributed network of noise-resistant neurons in the central auditory system

**DOI:** 10.1101/2020.06.10.143644

**Authors:** S. Souffi, C. Lorenzi, C. Huetz, J.-M. Edeline

## Abstract

Background noise strongly penalizes auditory perception of speech in humans or vocalizations in animals. Despite this, auditory neurons successfully detect and discriminate behaviorally salient sounds even when the signal-to-noise ratio is quite poor. Here, we collected neuronal recordings in cochlear nucleus, inferior colliculus, auditory thalamus, primary and secondary auditory cortex in response to vocalizations presented either against a stationary or a chorus noise. Using a clustering approach, we provide evidence that five behaviors exist at each level of the auditory system from neurons with high fidelity representations of the target, named target-specific neurons, mostly found in inferior colliculus and thalamus, to neurons with high fidelity representations of the noise, named masker-specific neurons mostly found in cochlear nucleus in stationary noise but in similar proportions in each structure in chorus noise. This indicates that the neural bases of auditory perception in noise rely on a distributed network along the auditory system.

## Introduction

In natural conditions, speech (in humans) and communication sounds (in animals) usually co-occur with many other competing acoustic signals. Both humans and animals exhibit remarkable abilities to reliably detect, process and discriminate communication sounds even when the signal-to-noise ratio (SNR) is quite low (Cherry, 1953; Gerhardt and Klump, 1988; Hulse et al., 1997). The auditory system has developed strategies to extract these behaviorally important signals mixed up with substantial amounts of noise. Over the last decade, many studies performed on different species have reported that the responses of auditory cortex neurons are quite resistant to various types of noises, even at low SNR (Narayan et al., 2007; Schneider and Woolley, 2013; Rabinowitz et al., 2013, Mesgarani et al., 2014; Ni et al., 2017; Beetz et al., 2018). Several hypotheses have been formulated to account for the high performance of auditory cortex neurons. For example, it was proposed that noise tolerance is correlated with adaptation to the stimulus statistics, which is more pronounced at the cortical than at the subcortical level (Rabinowitz et al., 2013). A dynamic model of synaptic depression combined with feedback gain normalization was also suggested as a potential mechanism for robust speech representation in the auditory cortex (Mesgarani et al., 2014). Alternatively, a simple feedforward inhibition circuit operating in a sparse coding scheme was viewed as a mechanism to explain background-invariant responses detected for a population of neurons in the secondary auditory cortex (Schneider and Woolley, 2013).

A recent study (Ni et al., 2017) reported that auditory cortex neurons can be assigned to categories depending upon their robustness to noise. More precisely, by testing the responses to conspecific vocalizations at different SNRs, this study described four types of responses classes (robust, balanced, insensitive and brittle) in the marmoset primary auditory cortex, and pointed out that depending upon the background noise, the same neuron can exhibit different response classes (Ni et al., 2017). The aim of the present study was to determine whether the subcortical auditory structures display similar proportions of these four types of response classes and whether the noise-type sensitivity is present at the subcortical level. We used exactly the same methodology as in Ni and colleagues (2017) to assign the neurons to a given response class: the Extraction Index (EI, initially defined by Schneider and Woolley, 2013) was computed at three SNRs (+10, 0 and −10 dB) and an unsupervised clustering approach (the K-means algorithm) was used to reveal groups of EI profiles. Based on this clustering approach performed on the whole auditory system, we define a typology of neuronal behaviors in noise based on five categories found at each stage, from target-ultraspecific and target-specific neurons showing a high fidelity representation of the target, to masker-specific neurons showing a high-fidelity representation of the noise, with two intermediary neuronal categories, one showing no preference either for the target or for the noise named non-specific neurons, and the other characterized by a sensitivity to the SNR named SNR dependent neurons.

Here, we present evidence demonstrating that the target-specific neurons are in higher proportions in inferior colliculus and thalamus in both noises, whereas the masker-specific neurons are found mostly in the cochlear nucleus in stationary noise but in similar proportions in each structure in a noise composed of a mixture of conspecific vocalizations that we will name “chorus noise”. We also provide evidence that the noise-type sensitivity - that is the ability to switch category from a given background noise to another - although present at each level of the auditory system in small proportions, is mostly detected in the inferior colliculus and the thalamus.

## Results

From a database of 2334 multi-unit recordings collected in the five investigated auditory structures, several criteria were used to include each neuronal recording in our analyses (see Table 1). A recording had to show significant responses to pure tones (see Methods section) and an evoked firing rate significantly above spontaneous firing rate (200 ms before each original vocalization) for at least one of the four original vocalizations (Fig. 1A illustrates their temporal envelopes and spectrograms). These four vocalizations were presented in quiet and embedded either in a vocalization-shaped stationary noise (Fig. 1B) or in a chorus noise (Fig. 1C) using three SNRs. We selected neurons showing responses at the three SNRs both in stationary and chorus noise in order to derive systematically six EI values for each neuronal recording. To determine a significance level of the EI value, we computed an EI_Surrogate_ value for each recording (see Methods section) and included only the recordings with at least one of the six EI values significantly higher than the EI_Surrogate_. Applying these criteria, we selected a total of 1267 recordings (Table 1): 389 neurons in the cochlear nucleus (CN), 339 neurons in the central nucleus of the inferior colliculus (CNIC), 198 neurons in auditory thalamus (MGv), 261 neurons in the primary auditory cortex (A1) and 80 neurons in a secondary auditory cortical area (VRB).

**Table 1.** Summary of the number of animals and number of selected recordings in each structure

|  | CN | Lemniscal pathway |  |  | Non-lemniscal pathway | Total |
| --- | --- | --- | --- | --- | --- | --- |
|  |  | CNIC | MGv | A1 | VRB |  |
| Number of animals | 10 | 11 | 10 | 11 | 5 | 47 |
| Number of recordings tested | 672 | 478 | 448 | 544 | 192 | 2334 |
| TFRP only | 560 | 421 | 285 | 455 | 126 | 1847 |
| TFRP and significant response to at least one vocalization | 499 | 386 | 262 | 354 | 95 | 1596 |
| Six EI values (for the six SNRs) | 488 | 368 | 224 | 320 | 95 | 1495 |
| One of the six EI values significantly higher than the $EI_{\text{Surrogate}}$ | 389 | 339 | 198 | 261 | 80 | 1267 |

**Figure 1.**
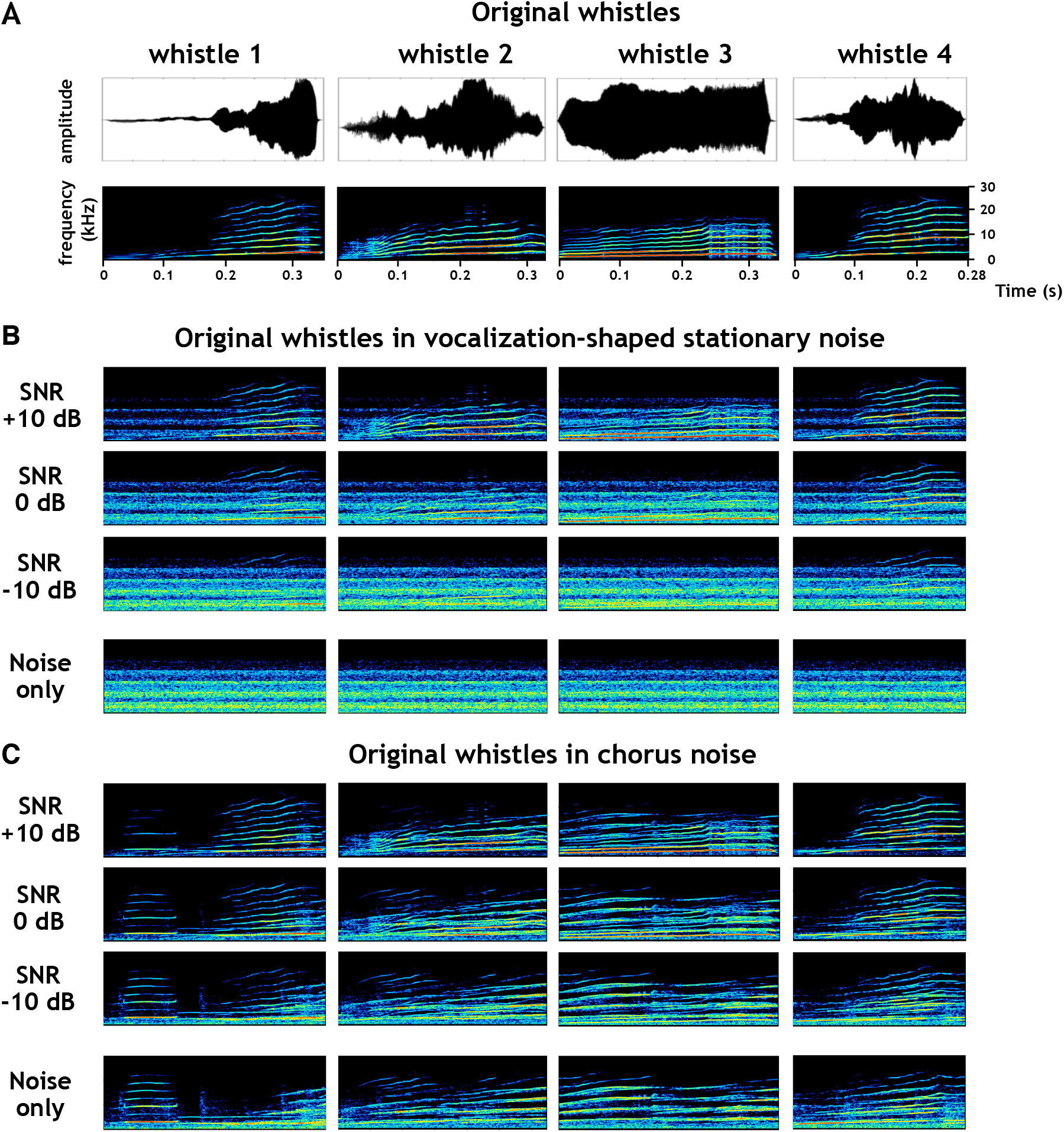
Original and noisy vocalizations. **A.** Waveforms (*top*) and spectrograms (*bottom*) of the four original whistles used in this study. **B-C.** Spectrograms of the four whistles in stationary (B) and chorus (C) noise at three SNRs (+10, 0 and −10 dB SPL, *from top to bottom*) and the noise only.

### Chorus noise impacted neuronal responses more than stationary noise at each stage of the auditory system

Figure 2A shows rasters for recordings collected in the five auditory structures in response to the original (in quiet) and masked vocalizations embedded in stationary and chorus noise. In all structures, the neuronal responses evoked by the four whistles progressively vanished as the SNR decreased from +10 to −10 dB. However, one can clearly detect that recordings obtained in CNIC and MGv still display clearly detectable responses at 0 dB SNR, even down to −10 dB for some vocalizations in CNIC.

**Figure 2.**
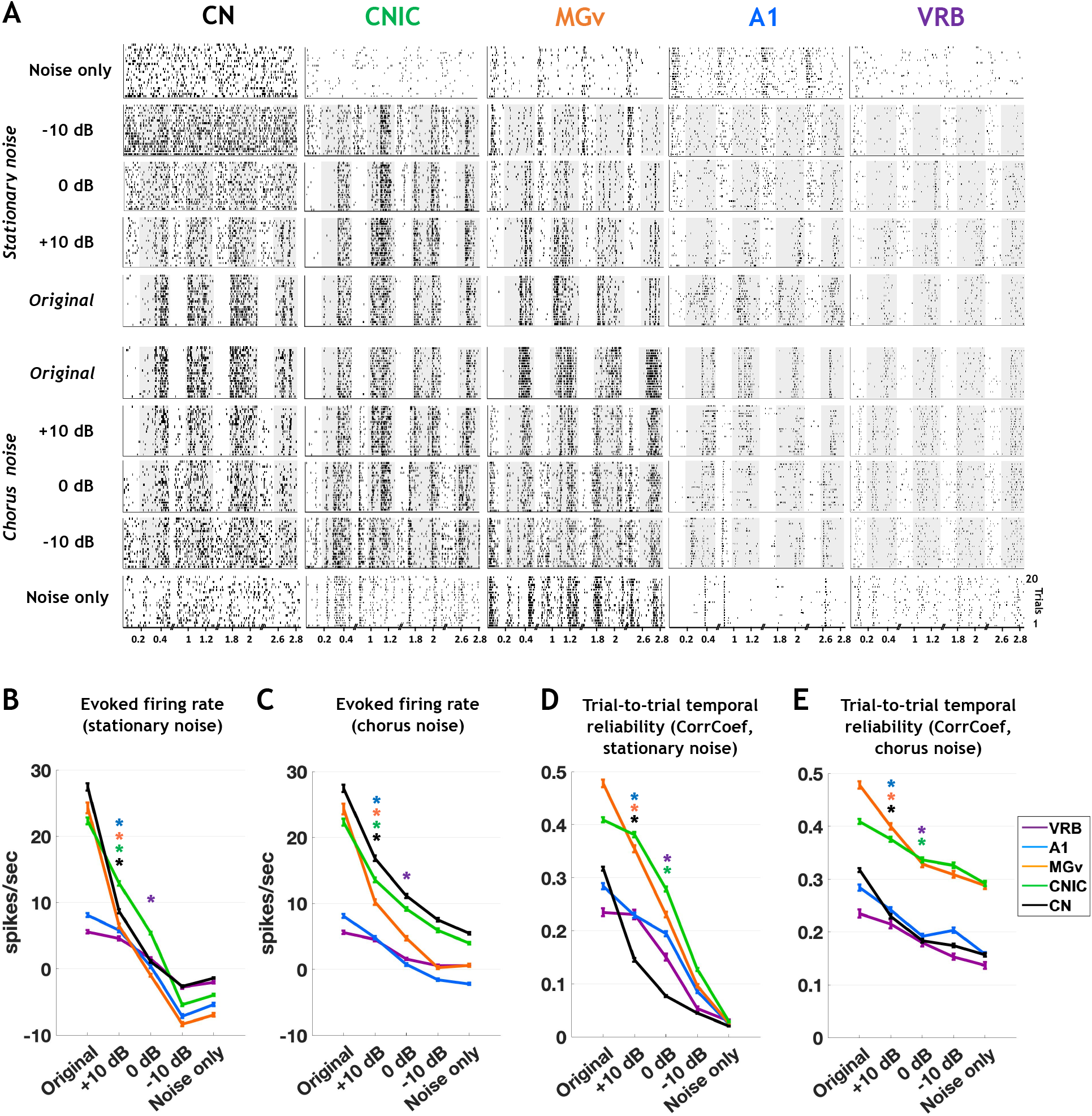
Noise strongly reduces the evoked firing rate and the temporal reliability but to a lesser extent for CNIC and MGv neurons. **A**. Raster plots of responses of the four original vocalizations, noisy vocalizations and noise alone in both noises recorded in CN, CNIC, MGv, A1 and VRB. The grey part of rasters corresponds to the evoked activity. **B-E.** The mean values (±SEM) represent the evoked firing rate (spikes/sec) (B, C) and the trial-to-trial temporal reliability represented by the CorrCoef value (D, E) obtained with original conditions, noisy vocalizations and noise alone in stationary and in chorus noise at three SNRs (+10, 0 and −10 dB SPL) in CN (in black), CNIC (in green), MGv (in orange), A1 (in blue) and VRB (in purple) (one-way ANOVA, P < 0.05; with post-hoc paired t tests, *P <0.05). The evoked firing rate corresponds to the total number of action potentials occurring during the presentation of the stimulus minus the spontaneous activity (200 ms before each acoustic stimulus).

Quantification of the evoked firing rate and the response trial-to-trial temporal reliability in stationary and chorus noise confirmed these observations (Fig. 2B-E). In both noises, the lower the SNR, the lower the evoked firing rate and the trial-to-trial reliability. More precisely, in both noises, the decrease in evoked firing rate was significant as early as the +10 dB SNR in all auditory structures except in VRB for which the decrease was significant at 0 dB SNR (Fig. 2B-2D; for the stationary noise (Fig. 2B): one-way ANOVA: F_CN(4,1944)_=315; F_CNIC(4,1694)_=265.5; F_MGv(4,989)_=174.9; F_A1(4,1304)_=95.8; F_VRB(4,399)_=40.8, p<0.001; with post-hoc paired t tests, p<0.001; for the chorus noise (Fig. 2D): one-way ANOVA: FCN(4,1944)=108.7; FCNIC(4,1694)=92.7; FMGv(4,989)=93.8; FA1(4,1304)=74.7; FVRB(4,399)=24.2, p<0.001; with post-hoc paired t tests, p<0.001). Similarly, in both noises, the trial-to-trial temporal reliability (quantified by the CorrCoef index) was significantly decreased as early as the +10 dB SNR in CN, MGv and A1 whereas in CNIC and VRB, the decrease was significant only at the 0 dB SNR (Fig. 2C-2E; for the stationary noise (Fig. 2C): one-way ANOVA: F_CN(4,1914)_=458.7; F_CNIC(4,1559)_=317.1; F_MGv(4,831)_=226.6; F_A1(4,1101)_=115.8; F_VRB(4,357)_=45.3, p<0.001; with post-hoc paired t tests, p<0.001; for the chorus noise (Fig. 2E): one-way ANOVA: F_CN(4,1916)_=58.9; F_CNIC(4,1614)_=16.6; F_MGv(4,929)_=28.4; F_A1(4,1096)_=18.8; F_VRB(4,365)_=6.5, p<0.001; with post-hoc paired t tests, p<0.001). Note that, on average, collicular and thalamic neurons showed higher temporal reliability than cochlear nucleus and cortical neurons both in quiet and in noise conditions (Fig. 2C-2E). Thus, in all structures, the two types of background noise decreased the firing rate and temporal reliability of neuronal responses. Even if the noise-induced changes for these two parameters were larger in subcortical structures compared to auditory cortex, the highest values remained in the inferior colliculus and thalamus. Neuronal responses were further investigated by quantifying the Extraction Index (EI, see Methods section; Schneider and Woolley, 2013; Ni et al., 2017) on our entire database, i.e. the 1267 recordings obtained in the five structures. For each neuron, this index compares the PSTH obtained at a given SNR with the PSTHs obtained with the original vocalizations and with the noise alone: the higher the EI value (close to 1), the more the responses are target-like. Conversely, the lower the EI value (close to −1), the more the responses are masker-like.

We found that the mean EI values were higher in the inferior colliculus and thalamus than in the cochlear nucleus and cortex, except in chorus noise at – 10 dB SNR, which strongly impacted all neuronal responses at each stage (Fig. 3). Figure 3A displays examples of rasters illustrating a case of EI > 0 (left panel) and a case of EI < 0 (right panel). Figure 3 also presents the EI values in chorus noise as a function of EI values in stationary noise at the +10, 0 and −10 dB SNR for all the recordings (Fig. 3B-D). The increase in the number of dots in the bottom left quadrant indicates that for most of the neurons, the EI decreased from +10 dB to 0 dB SNR, an effect which is further accentuated at −10 dB SNR. In addition, more pronounced effects of the chorus noise were observed as early as the 0 dB SNR in each auditory structure as indicated by the large number of dots below the diagonal lines. Examination of these scattergrams also indicates that the EI value distributions differed between structures (represented by a color code). Statistical analyses revealed that on average, EI values decreased when the SNR decreased in all structures and for both types of noises (for the stationary noise (Fig. 3E): one-way ANOVA: F_CN(2,1166)_=171.1; F_CNIC(2,1016)_=176.3; F_MGv(2,593)_=77.6; F_A1(2,782)_=37.6; F_VRB(2,239)_=14.3, p<0.001; with post-hoc paired t tests, p<0.001; for the chorus noise (Fig. 3F): one-way ANOVA: FCN(2,1166)=394.4; FCNIC(2,1016)=361.0; FMGv(2,593)=275.3; FA1(2,782)=104.5 FVRB(2,239)=47.6, p<0.001; with post-hoc paired t tests, p<0.001).

**Figure 3.**
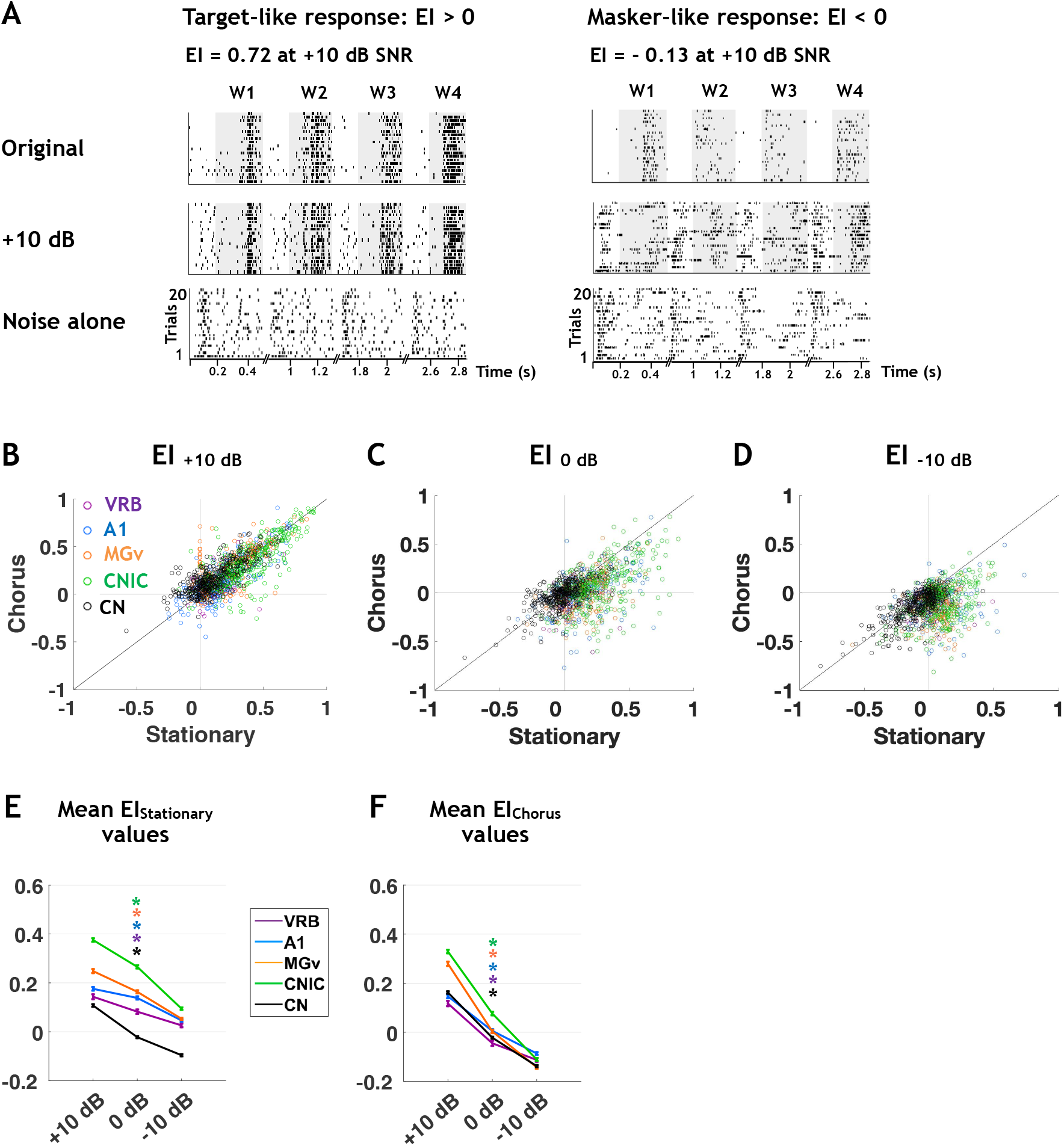
The decrease of EI values in each auditory structure is more pronounced in chorus noise than in stationary noise. **A.** Examples of neuronal responses in stationary noise with values of EI > 0 corresponding to a vocalization-like response (left) and EI < 0 corresponding to a noise-like response (right). Top panels shows the responses to the original vocalizations, the middle panels the responses to vocalizations at the +10 dB SNR and the bottom panels the responses to stationary noise alone. **B-F.** Scattergrams showing the SNR effect on EI values in chorus noise as a function of EI values in stationary noise at +10, 0 and −10 dB SNR (B-D) and mean EI values (±SEM) for the three SNRs (E-F) obtained in CN (in black), CNIC (in green), MGv (in orange), A1 (in blue) and VRB (in purple) in stationary (E) and chorus noise (F).

Thus, in all structures, both noises alter the evoked responses promoting masker-like responses, and the chorus noise promoted a higher number of masker-like responses than the stationary noise.

We next aimed at characterizing categories of neurons that display a particular behavior in noise in relation to fidelity of neural representation either of the target or of the noise. Therefore, a clustering analysis was performed on the entire database, i.e. the 1267 recordings obtained in the five structures, separately for the stationary and for the chorus noise.

### Five distinct groups of neurons exist at each stage of the auditory system but in different proportions

We initially aimed at determining whether the four categories of cortical neurons (robust, balanced, insensitive and brittle) described by Ni and colleagues (2017) across several SNRs can also be found at each stage of the auditory system. Analyzing our whole database with the same clustering method and the same criteria (elbow method) as in Ni and colleagues (2017) led us to consider either five or six clusters (Fig. 4A) in both noises. When six clusters were defined, two of them displayed very similar behaviors with only slight differences in EI values in each noise type (see Fig. 4B-C), which urged us to consider only five clusters. Compared to the four categories of Ni and colleagues (2017), we added one new category which represents an attenuated version of their robust neurons. These neurons represent in fact a large proportion of our database (25% and 18% in the stationary and chorus noise) and cannot be neglected.

We also opted for cluster names using more symmetric terms. The neurons keeping a high-fidelity representation of the vocalizations despite the presence of noise will be called the target-ultraspecific or target-specific neurons, those keeping a high-fidelity representation of the noise across the SNRs will be called masker-specific neurons and those showing no preference will be called non-specific neurons. One last category of neurons is characterized by sensitivity to the signal-to-noise ratio and therefore will be called SNR-dependent. This change in cluster names gives an equivalent role to target-specific and noise-sensitive neurons, named here masker-specific neurons, since in some ethological conditions they could play a functional role as important as the target-specific neurons.

**Figure 4.**
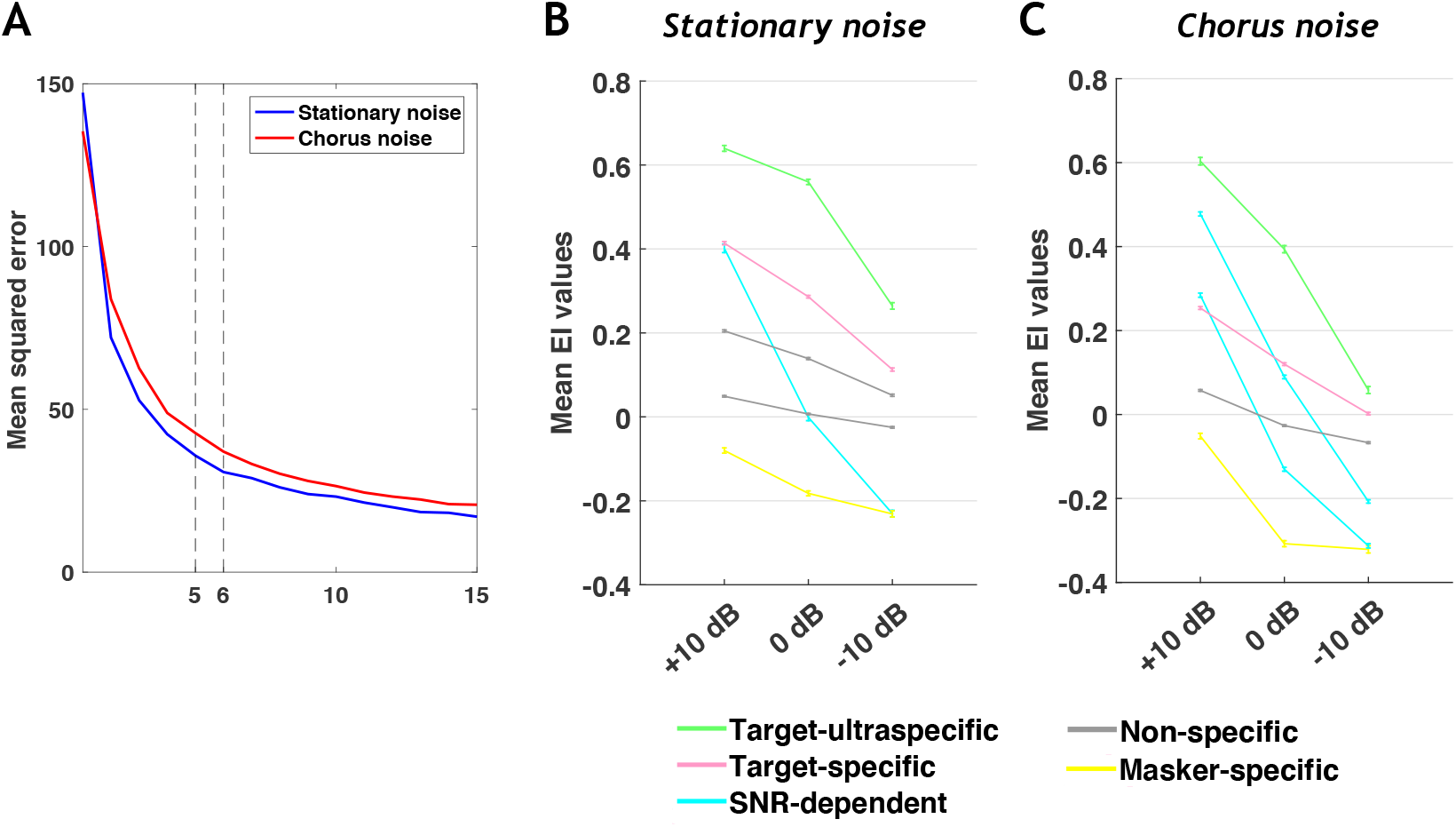
The choice of five clusters is optimal to reveal the different behaviors in both noises. **A.** Mean square error of EI profile clustering as a function of the number of clusters using the K-means algorithm for the stationary and chorus noise. **B-C.** Population average EI profile (±SEM) of each cluster when considering six clusters to separate the data in the stationary noise (B) and in the chorus noise (C). Note that in both noises, two clusters have equivalent mean EI profile (i.e. the same EI evolution across the three SNRs) and this led us to consider only five clusters in the results section (Figure 5).

Figure 5 presents the five clusters derived from the whole data set (from cochlear nucleus to secondary auditory cortical field) across the three SNRs and in the two noise conditions. Figures 5A and 5F present the EI values of all neurons (with a color code from blue to red when progressing from low to high EI values) and the color bars, on the right side, delineate the five clusters. This color code was used for the 3D representations of the five clusters in the stationary (Fig. 5B) and chorus (Fig. 5G) noise. Figure 5C shows the mean EI values in stationary noise for these five clusters across the three SNRs and the percentage of neurons in each cluster is displayed in figure 5D. Approximately 10% of the neurons are target-ultraspecific characterized, on average, by EI values greater than 0.5 at +10 and 0 dB SNRs. More than 25% are target-specific characterized, on average, by EI values greater than 0.2 at +10 and 0 dB SNRs (Fig. 5C). About 5% of the neurons are SNR-dependent and more than 40% of the total population has EI values around 0 at all SNRs which corresponds to the non-specific neurons. More than 10% of the auditory neurons have negative EI values at the three SNRs and correspond to masker-specific neurons. Figures 5H and 5I show the mean EI values for these five clusters in the chorus noise with, roughly, similar proportions of the five clusters as in the stationary noise. However, in the chorus noise there was a decrease in the proportions of neurons in the target-ultraspecific (from 10% to 7.5%) and target-specific (from 27% to 20%) clusters associated with an increase in the proportion of SNR-dependent neurons (from 6.5% to 19.5%), whereas the proportion of neurons in the non-specific cluster remained similar (42-39.5%). Note also that in the chorus noise, the two clusters of target-specific neurons showed, on average, lower EI values at the 0 dB SNR than in stationary noise (compared Fig. 5C and Fig. 5H). Based upon these quantifications, it is clear that, in the entire auditory system, the chorus noise impacted more the neuronal responses than the stationary noise.

**Figure 5.**
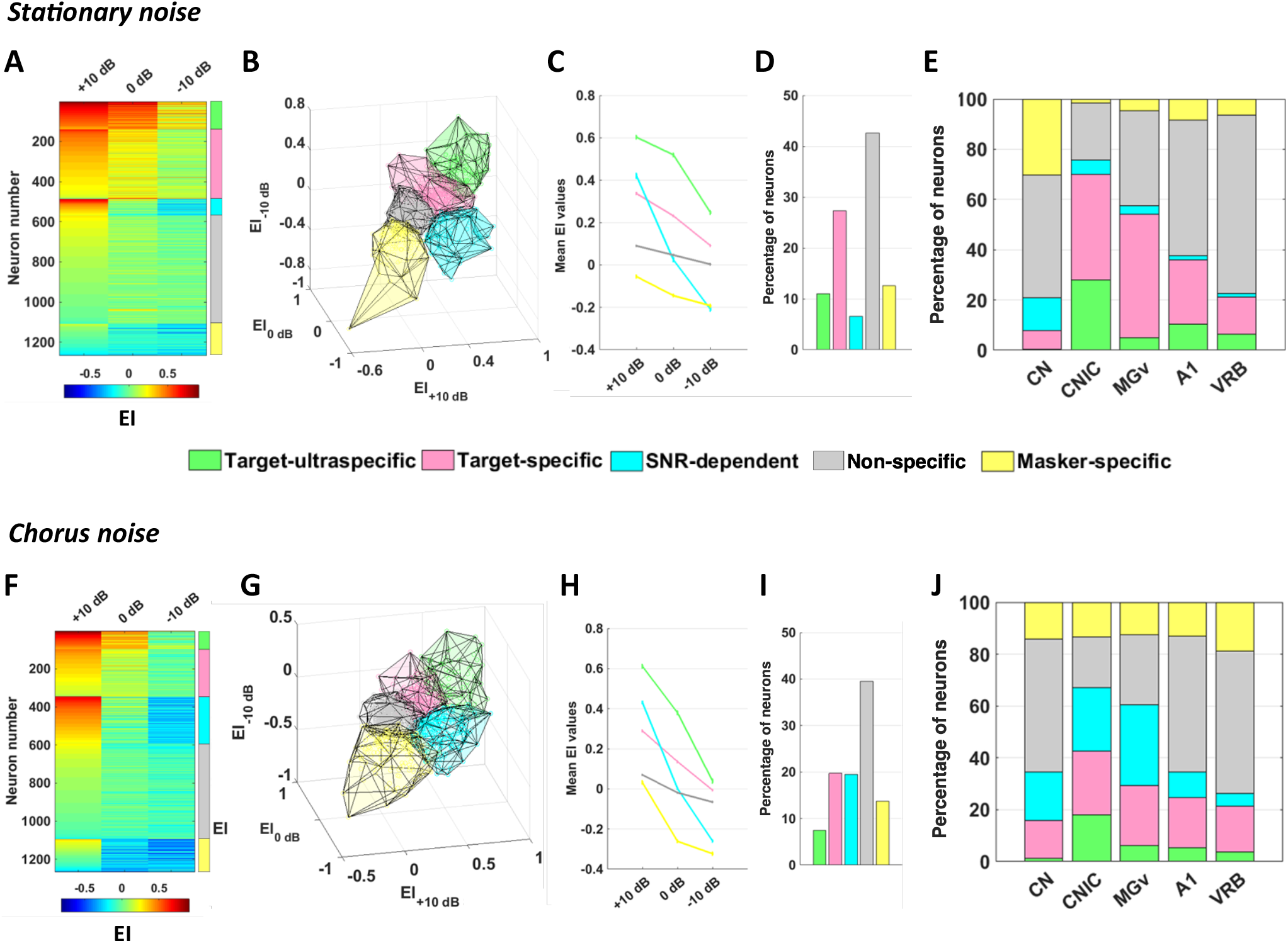
Five different groups of auditory neurons in noise. **A.** Each row corresponds to the EI profile of a given neuronal recording obtained in the five auditory structures in stationary noise with a color code from blue to red when progressing from low to high EI values. On the right, five stacked colors delineate the cluster identity for the five groups of neurons. The target-ultraspecific group is in green, the target-specific group in pink, the SNR-dependent group in turquoise, the non-specific group in gray and the masker-specific group in yellow. **B-E** 3D representation of the five clusters in stationary noise (B), mean EI values of the five clusters (C), relative proportions of each cluster in stationary noise (D) and proportion of each cluster in the five auditory structures from CN to VRB (E). **F-J.** Same representations as in A, B, C, D and E for the chorus noise. **Figure supplement 1.** Five neuronal behaviors in noise are found at each stage of auditory system.

**Figure 5 - figure supplement 1.**
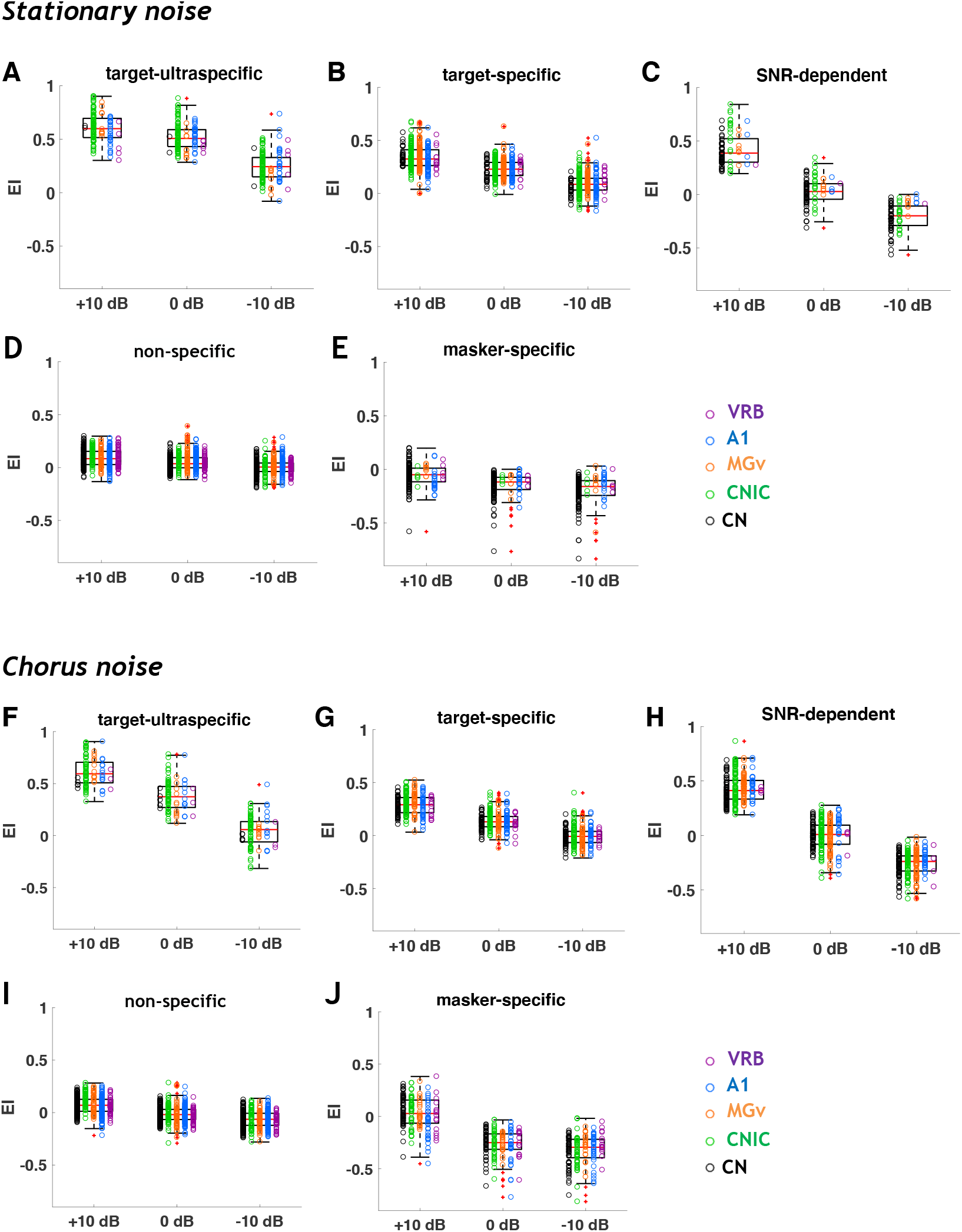
Five neuronal behaviors in noise are found at each stage of auditory system. **A-E.** The boxplots show all EI values obtained for the three SNRs tested (+10, 0 and −10 dB) in stationary noise for each auditory structure (CN, (in black), CNIC (in green), MGv (in orange), A1 (in blue) and VRB (in purple)) depending on the cluster assigned. **F-J.** Same representations as in A, B, C, D and E for the chorus noise.

Next, we determined the proportions of each cluster in a given structure. In stationary noise, target-ultraspecific and target-specific neurons were mostly present in the inferior colliculus and thalamus, while the three other groups of neurons classified as SNR-dependent, non-specific, and masker-specific were mostly present in the cochlear nucleus and in the two cortical fields. For each auditory structure, the percentage of neurons from each cluster is presented in the stationary noise (Fig. 5E) and in the chorus noise (Fig. 5J). Statistical analyses confirmed that the proportion of the different clusters differed in the IC and MGv compared with the three other structures (all Chi-Square; p<0.001). In chorus noise, the proportion of target-ultraspecific and target-specific neurons decreased in all structures but they remained in higher proportions in IC and MGv than in CN and in cortex. In the CN, there was also an increase in the proportion of target-specific neurons (from 7 to 14.5%). Statistical analyses confirmed that, in the chorus noise too, the proportion of the different clusters differed in the IC and MGv compared with the three other structures (all Chi-Square; p<0.001).

Plotting all EI values for the five clusters showed a good homogeneity across structures within each cluster type either in stationary or in chorus noise (Figure 5 - figure supplement 1), implying that a cortical neuron assigned to a particular cluster has the same behavior in noise than a subcortical neuron.

Therefore, in both noises, the neurons with a high fidelity representation of the target were mostly present in the inferior colliculus and thalamus. The non-specific neurons showing no preference either for the target or the noise were found in majority in the cochlear nucleus and in the auditory cortex. The SNR-dependent neurons represented a small fraction of neurons in stationary noise but were more numerous in the chorus noise. One interesting feature is that, in both types of noise, the proportion of these SNR-dependent neurons decreases progressively as one ascends in the auditory system. Finally, the neurons with a high fidelity representation of the noise were mostly localized in the cochlear nucleus in the stationary noise but were in an equivalent proportion in all structures (between 12.5% and 19%) in the chorus noise.

### Collicular and thalamic neurons are most sensitive to the type of background noise

In the marmoset auditory cortex, Ni and colleagues (2017) have pointed out that the neuronal behavior in noise can be context-dependent: the behavior of a given neuron in a particular noise does not predict its behavior in another noise. Is this a property that characterizes cortical neurons, or is it a property that exists at all levels of the auditory system?

Overall, we found that the neuronal response behaviors at all levels of the auditory pathway were partly, but not completely, preserved in different types of noise (Fig. 6A). On the whole database of 1267 recordings, about 50% of the target-ultraspecific and 40% of the target-specific neurons in the stationary noise remained so in the chorus noise; most of the SNR-dependent (73.5%) and non-specific neurons (65.5%) in the stationary noise remained also in the same category in the chorus noise. Only the masker-specific neurons tended to switch category to mostly became non-specific neurons. Considering the proportions of neurons in each category in stationary noise, as indicated on the x-axis of figure 6A, around 45% of all recordings switched category in chorus noise (see Fig. 6B). Figure 6B shows that the target-ultraspecific, target-specific and masker-specific neurons were the three categories with the highest percentages of cluster changes (χ^2^ p<0.001). We used a bootstrap procedure to have a better estimation of the percentage of cluster changes (see Methods section). Briefly, for each recording, and from the 20 trials obtained for each stimulus, we resampled 20 trials (allowing repetitions), recomputed the Extraction Index and reallocated each resampled recording in the closest cluster. This entire procedure was performed 100 times for each recording. We assumed that in a given type of noise, a recording could change category because of its response variability and/or because it was located very close to a border between two clusters, independently to the change in noise type. With the resampled data obtained from the bootstrap procedure, we determined, for each cluster type (Fig. 6B), and each structure (Fig. 6C), the percentage of cluster change averaged for the two types of noise.

**Figure 6.**
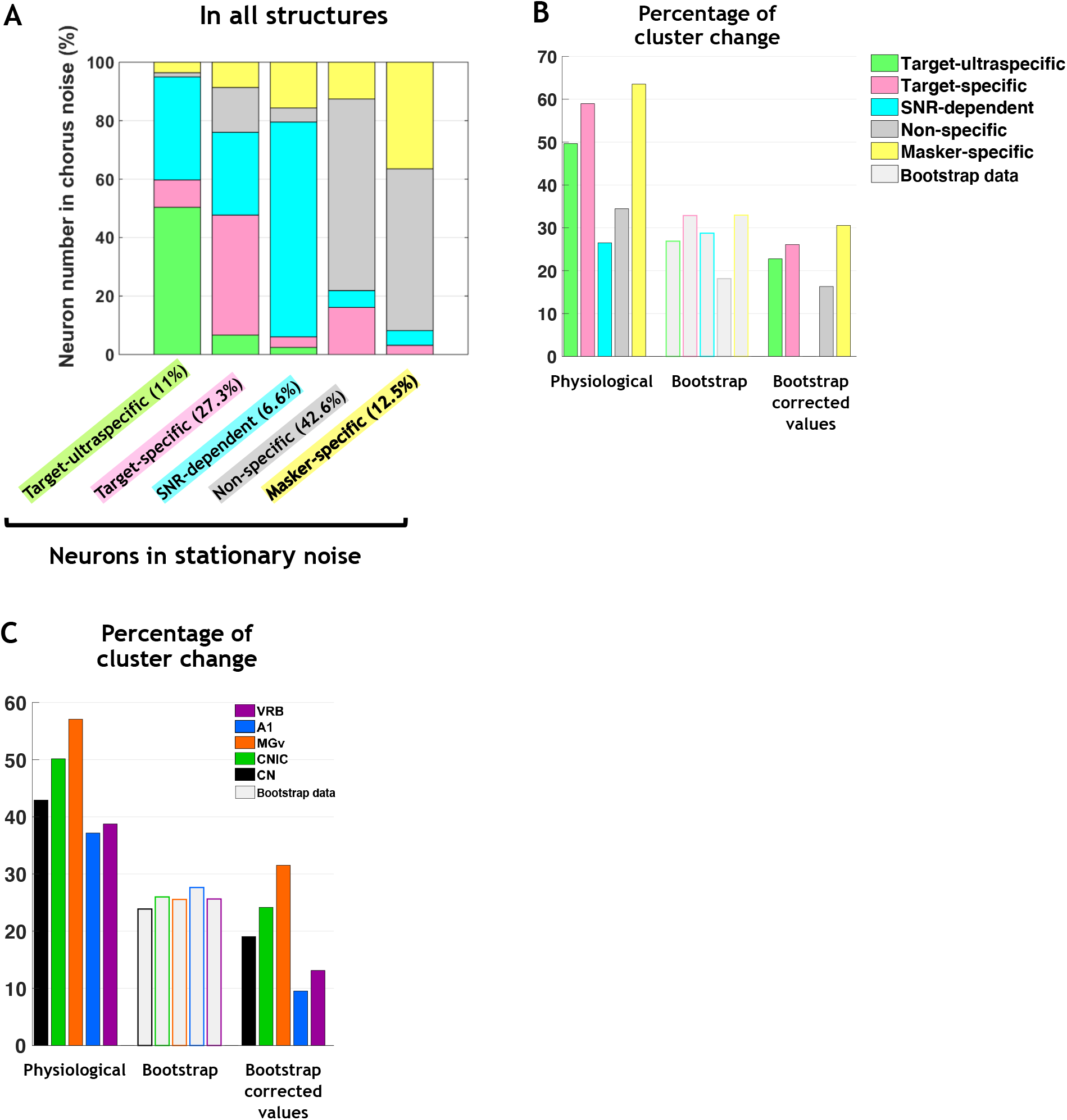
Clustering change from one background noise to another is found at each stage of the auditory system. **A.** Percentage of neurons in a given cluster in chorus noise depending on the cluster originally assigned in the stationary noise. The abscissa indicate the cluster identity in the stationary noise and the ordinates represent the cluster identity in the chorus noise. For example, target-ultraspecific neurons in stationary noise are also 50% target-ultraspecific in chorus noise but 10% are reclassified as target-specific, 35% SNR-dependent, 1.5% non-specific and 3.5% masker-specific. **B.** Percentage of neurons changing of cluster from the stationary noise to the chorus noise for each cluster computed based on physiological and bootstrapped data (grey bars). The corrected-bootstrap values correspond to the subtraction of the bootstrapped values from the physiological values. **C.** Mean percentage of cluster change from the stationary noise to the chorus noise in VRB, (in purple), A1 (in blue), MGv (in orange), CNIC (in green) and CN (in black). The grey bars represents the percentage of cluster change computed based on bootstrap analysis (see Method section). **Figure supplement 1.** Bootstrapped data.

For each cluster type, except for the non-specific cluster, around 25% of resampled data switched category, which is indicated by the grey section (Fig. 6B). For the non-specific cluster, less than 20% changed category with the resampled data (Fig. 6B). When subtracting these bootstrapped percentages, we obtained the bootstrap-corrected values of the percentage of cluster change in each cluster, which dropped the percentages of cluster changes obtained with physiological data to only 16-30% for the target-ultraspecific, target-specific, non-specific and masker-specific neurons. For the SNR-dependent neurons, the cluster change observed with physiological data is absent when subtracting the bootstrapped percentage obtained for this cluster type suggesting that the SNR-dependent neurons changed category very little or not at all in chorus noise.

We performed the same analysis for each auditory structure (Figure 6 - figure supplement 1, Fig. 6C). On average, between 21 and 31 out of 100 bootstrapped data per recording changed cluster in cortical and subcortical structures in both noises (Figure 6 - figure supplement 1). Thus, with only the resampled data, there is, on average, an important fraction of the recordings changed cluster in each structure suggesting that the response variability and/or the proximity of the borders between two clusters induced cluster changes. Figure 6C presents the mean percentage of cluster change obtained from the physiological data for each auditory structure: for the two cortical areas about 38% of the neurons changed categories, it was 57% in the MGv, 50% in the inferior colliculus and 43% in the cochlear nucleus. With bootstrapped data, in all auditory structures, between 20 to 30% of the neurons changed category, which is indicated by the grey section. When subtracting these bootstrapped percentages, we obtained the bootstrap-corrected values of the percentages of cluster changes in each structure, which dropped the percentage of cluster change obtained with physiological data to only 10-30% (Fig. 6C). The percentage of neurons changing category was higher in the inferior colliculus and thalamus (31-22%) than in the cochlear nucleus and the auditory cortex (14% on average; Fig. 6C). Therefore, the noise-type sensitivity is present at each stage of the auditory system but represents small proportions. Furthermore, the inferior colliculus and the thalamus had the most noise-type sensitive neurons, the fewest were found in the auditory cortex.

**Figure 6 - figure supplement 1.**
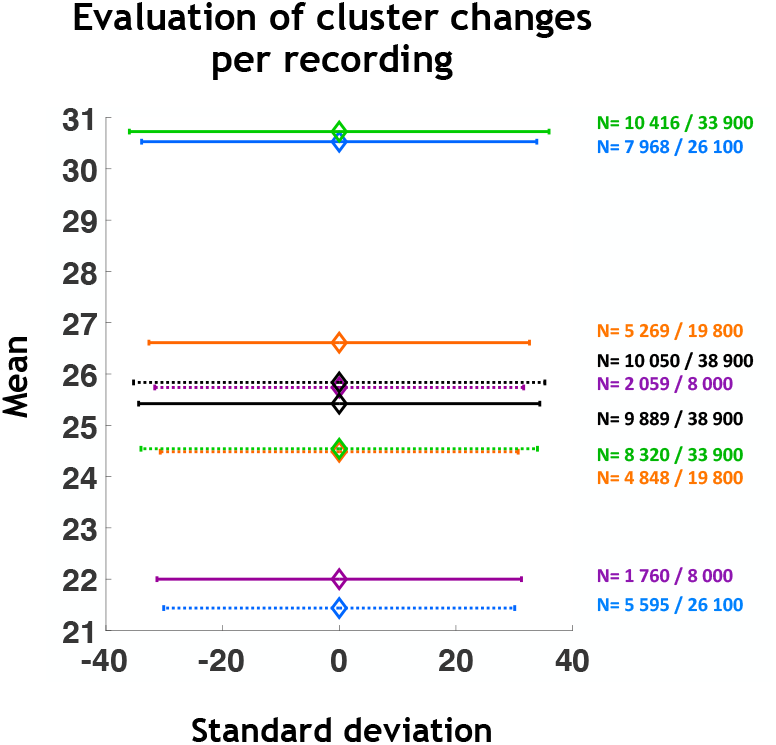
Bootstrapped data. Means and standard deviations of cluster changes per recording obtained with bootstrapped data in stationary noise (solid lines) and chorus noise (dotted lines) for each auditory structure. The numbers to the right of the figure indicate the cumulative bootstrapped data. For example, in thalamus and in stationary noise (orange solid lines), out of 19 800 bootstrap data, 5 269 (26.6%) changed cluster from stationary noise to the chorus noise.

One of the questions that should be addressed is whether or not the robustness to noise detected here is correlated to the response characteristics obtained with the original vocalizations in quiet. For example, one can envision that the EI value (and its evolution across the three SNRs) is related either to the strength or temporal reliability of the responses, or the ability to discriminate the four original vocalizations. Therefore, in both noises, we looked for potential correlations between the EI values and the response parameters to the original vocalizations. We focused only on the neurons exhibiting the same neuronal behavior in the two noises (n_Total_=685, n_CN_=222, n_CNIC_=169, n_MGv_=81, n_A1_=164, n_VRB_=49) to look for correlations between stable EI values and other parameters obtained in quiet. For these neurons, the evoked firing rate was significantly higher in the subcortical structures than in the cortex (unpaired t-test, lowest p-value p<0.001; Fig. 7A). The CorrCoef values were significantly higher in CNIC and MGv compared to A1 and VRB (Fig. 7B), and the MI_Individual_ values obtained at the subcortical level were also significantly higher than at the cortical level (unpaired t-test, highest p<0.001 between the cortex and the subcortical structures; Fig. 7C).

**Figure 7.**
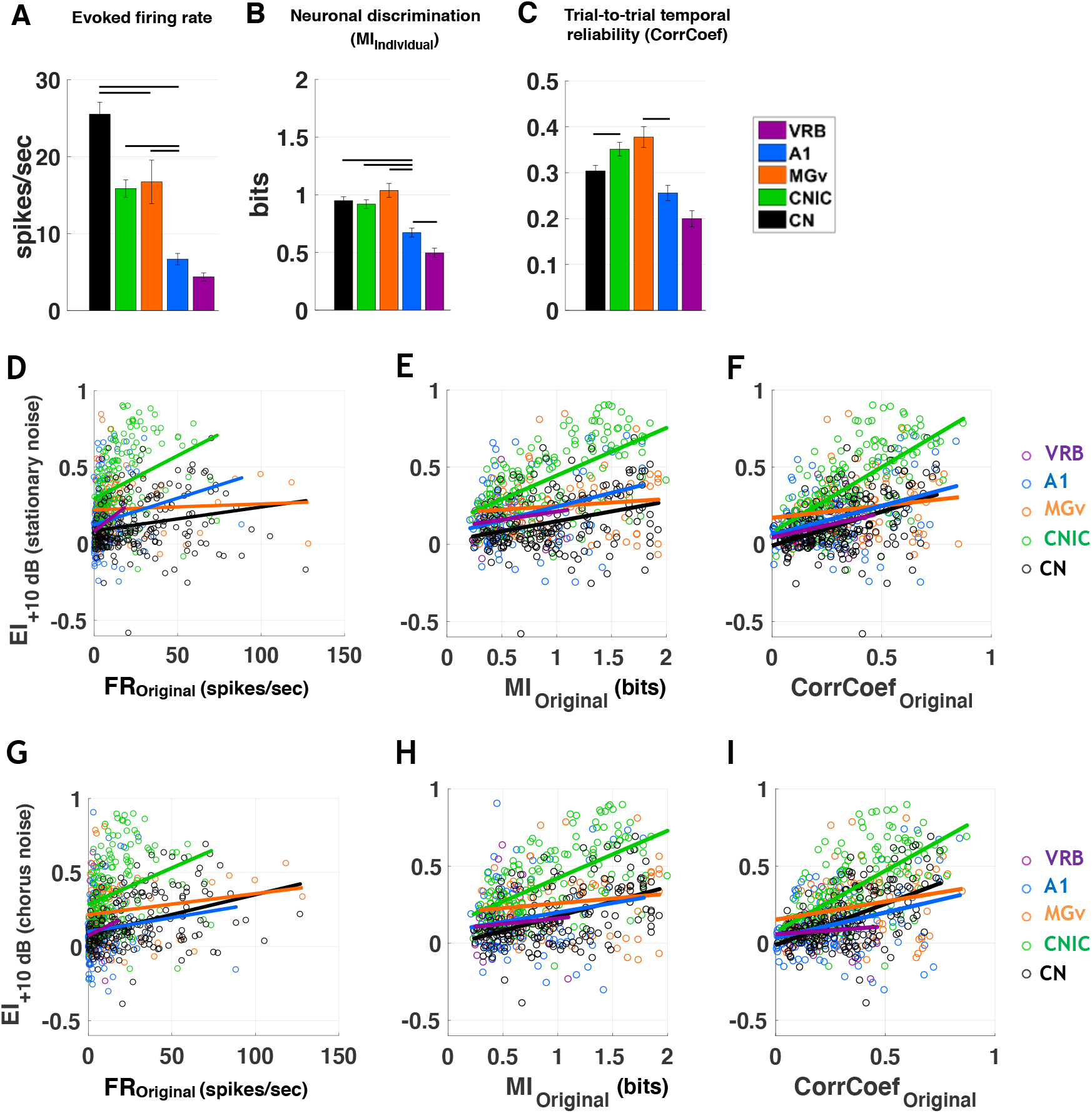
The trial-to-trial temporal reliability of responses in original conditions is correlated with the capacity of neurons to resist in noise. **A-C.** The bar graphs show the mean values of (**A**) the evoked firing rate (spikes/sec), the neuronal discrimination assessed by the mutual information (MI) computed at the level of the individual recording (MI_Individual_, bits) (**B**) and the trial-to-trial temporal reliability quantified by the CorrCoef value (**C**) obtained with the four original vocalizations for only the neurons that remained in the same category in both noises in CN (in black), CNIC (in green), MGv (in orange), A1 (in blue) and VRB (in purple). The evoked firing rate corresponds to the total number of action potentials occurring during the presentation of the stimulus minus spontaneous activity (200 ms before each acoustic stimulus). In each structure, error bars represent the SEM of the mean values and black lines represent significant differences between the mean values (unpaired t test, p<0.05). The evoked firing rate decreases from the CN to VRB but both the trial-to-trial temporal reliability (CorrCoef) and the discrimination performance (MI) reach a maximal value in subcortical structures. **D-F.** Scattergrams of the EI values obtained at the +10 dB SNR in stationary noise as a function of the evoked firing rate (in spikes/sec), mutual information (MI, bits) and CorrCoef values obtained with the original vocalizations based on neuronal responses recorded in CN, CNIC, MGv, A1 and VRB for only the neurons that remained in the same category in both noises. **G-I.** Same representations as in D, E, F and d for the EI values obtained at the +10 dB SNR in chorus noise.

Generally, both for the stationary noise (Fig. 7D-F) and for the chorus noise (Fig. 7G-I), the strongest correlations were found between the EI values and the CorrCoef values (Fig. 7F, 7I; for example r_CNIC_ = 0.65; r_CN_ = 0.45; r_A1_ = 0.41 for the stationary noise), whereas much weaker correlations (if any) were found with the evoked firing rate (Fig. 7D, 7G; for example r_CNIC_ = 0.34; r_CN_ = 0.21; r_A1_ = 0.17 for the stationary noise) or with the MI values (Fig. 7E, 7H; for example r_CNIC_ = 0.61; r_CN_ = 0.33; r_A1_ = 0.30 for the stationary noise). Except in one case, the same significant correlations were found in the chorus noise (see Table 2). Note that the strongest correlations between the EI and the CorrCoef values were obtained in the inferior colliculus (0.65 and 0.63 in stationary and chorus noise respectively). These results suggest that the trial-to-trial temporal reliability is the factor the more correlated with the robustness to noise, especially in the inferior colliculus, the auditory structure where we detected the largest number of target-specific neurons.

**Table 2.** Correlations values between the EI index obtained in both noises and the different parameters quantifying neuronal responses in quiet. (*) indicates significant correlations (p<0.05).

|  | Correlations with the EI index (r) |  |  |  |  |  |
| --- | --- | --- | --- | --- | --- | --- |
|  | Evoked FR |  | MI |  | CorrCoef |  |
|  | Stationary | Chorus | Stationary | Chorus | Stationary | Chorus |
| CN | 0.21* | 0.36* | 0.33* | 0.48* | 0.45* | 0.55* |
| CNIC | 0.34* | 0.30* | 0.61* | 0.61* | 0.65* | 0.63* |
| MGv | 0.05 | 0.18 | 0.13 | 0.16 | 0.17 | 0.24 |
| A1 | 0.17* | 0.09 | 0.30* | 0.20* | 0.41* | 0.34* |
| VRB | 0.23 | 0.13 | 0.16 | 0.08 | 0.32* | 0.09 |

## Discussion

Here, we demonstrate that the processing of noisy vocalizations by neurons in the entire auditory system can be described by a limited number of neuronal behaviors found in different proportions depending on the auditory structure and the type of the masking noise. Target-specific neurons were detected at each level of the auditory system but were in higher proportions at the collicular and thalamic level. For these neurons, the trial-to-trial temporal reliability of the responses in quiet is correlated to the robustness of responses in noise. In terms of proportions, the highest fidelity representation of the target or noise was found at the subcortical level whereas at the cortical level, the majority of neuronal responses showed no preference for the target or the noise suggesting that cortical neurons are more independent of the spectro-temporal characteristics of the noisy vocalizations. At each stage of the auditory system, neurons sensitive to the type of noise were observed in small proportions, mostly in inferior colliculus and thalamus.

### Robustness to noise in the auditory system: a localized vs. distributed property?

In the A1 of awake marmosets, Ni and colleagues (2017) found about 20-30% of robust neurons (depending on the vocalization), called here target-specific neurons. In our cortical data, when pooling together the target-ultraspecific and target-specific neurons, we obtained about the same proportions as in the marmoset A1 (33%). In the bird auditory system, Schneider and Woolley (2013) described the emergence of noise-invariant responses for a subset of cells (the broad spike cells) of a secondary auditory area (area NCM), whereas upstream neurons (IC and A1 neurons in their study) represent vocalizations with dense and background-corrupted responses. They suggest that the sparse coding scheme operating within NCM allows the emergence of this noise-invariant representation. In our study (and in the mammalian A1 in general), a sparse representation already exists as early as A1 (see the rasters in Fig. 2, see also Hromádka et al., 2008) allowing target-ultraspecific and target-specific neurons to be present in about the same proportions in A1 and the secondary area VRB.

Noise-invariant representations were also reported in A1 of anesthetized ferrets (Rabinowitz et al., 2013). This study suggested a progressive emergence of noise-invariant responses from the auditory nerve to IC and to A1, and proposed the adaptation to the noise statistics as a key mechanism to account for the noise-invariant representation in A1. However, a large fraction of their IC neurons also showed responses that were independent of the background noise (see their Fig. 2C) and adapted to the stimulus statistics (see their Fig. 4B), suggesting that the behavior of IC and A1 neurons did not fundamentally differ in their adaptation to noise statistics.

Based upon the proportion of target-ultraspecific and target-specific neurons, it seems that the robustness to noise peaks in CNIC, with the MGv neurons being at the intermediate level between IC and A1 (Figures 5E, 5J). In fact, our results point out an abrupt change from a prominent noise-sensitivity in CN to a prominent noise-robustness in IC, which means that this robustness is generated by neural computation taking place in the central auditory system. Whether this is an intrinsic property emerging *de novo* in the IC or whether this property emerges as a consequence of cortical feedbacks (Malmierca and Ryugo, 2011) which can promote behavioral plasticity (Bajo et al., 2010; Robinson et al., 2016) remains to be determined. Nonetheless, several studies have clearly demonstrated that IC neurons adapt to the stimulus statistics. First, adaptations of IC neurons to the average stimulus intensity, stimulus variance and bimodality that has already been described with a temporal decay of about 160 ms at 75 dB (Dean et al., 2005; 2008). Second, adaptation to the noise statistics shifted the temporal modulation function (TMF) of IC neurons to slower modulations, sometimes transforming band-pass TMF to low pass TMF in about 200 ms of noise presentation (Lesica and Groethe, 2008).

Our results do not indicate a progressive evolution from sensitivity to robustness to noise along the central auditory system. However, based upon the proportion of SNR-dependent neurons, one interesting result is that, in both types of noise, these neurons decreased progressively as one ascends in the auditory system which is in line with the idea of a progressive construction of an invariant representation of acoustic signals in noise described by Rabinowitz and colleagues (2013).

We also showed that the higher the trial-to-trial temporal reliability of the responses in quiet, the higher the robustness of the neurons in noise, especially in the IC where we detected the highest proportion of target-specific neurons. In fact, it was previously reported that the firing rate and the temporal reliability of IC neurons decreased when vocalizations were presented in natural stationary noise, but they were still efficiently detected target stimuli in noise (Lesica and Groethe, 2008). Together, this suggests that the more temporally precise are the synaptic inputs converging on a particular neuron, the more the responses of this neuron are robust in background noise.

The subcortical robustness to noise described here might be surprising given that numerous studies have pointed out the robustness of cortical representations in noise. However, Las and colleagues (2005) reported that A1, MGB and IC neurons can detect low-intensity target tones in a louder fluctuating masking noise and display the so-called “phase-locking suppression”, that is the interruption of phase-locking to the temporal envelope of background noise. The last result indicates that, as cortical neurons, both IC and MGB neurons have the ability to detect low-intensity target sounds in louder background noise (even at −15 or −35 dB SNR) and the robustness of some of our subcortical neurons may stem from this ability to detect target vocalizations even at SNR as low as the −10 dB SNR.

Robust perception of target sounds probably also requires a robust representation of competing sounds (here, masking noise). This can be the functional role of the masker-specific neurons, which are potentially crucial to determine the characteristics of the noise type and to provide an accurate representation of it within the auditory stream reaching our ears at any time. They were detected here, in higher proportion in the CN in stationary noise, but they became more numerous and in equivalent proportion in all structures in chorus noise. Therefore, the noise representation can be based upon the neuronal activity in the cochlear nucleus in stationary noise, whereas this representation can be more distributed in the chorus noise potentially because some of the target-ultraspecific or the target-specific neurons in stationary noise became masked-specific neurons in chorus noise, due to its spectrotemporal acoustic richness.

In our results, the five categories rather form a continuum with no clear boundaries between clusters, which inevitably led us to « impose » the clustering. Nonetheless, despite the lack of precise boundaries, 75% of the neurons remained assigned to the same cluster when the bootstrap procedure was performed (Fig. 6B-C), suggesting a relatively good reliability of the classification. Also, these five categories do represent distinct neuronal behaviors in the two types of noise, which have been previously described at the cortical level in awake marmoset (Ni et al., 2017). Here, in the auditory cortex of anesthetized animals, we found these same global behaviors, and our results show that these categories also exist at the subcortical level. Thus, the cortical representation of noisy signals by different neuronal categories characterized either by the preference of the target, the noise, a sensitivity to SNR or an absence of these three acoustic features, is independent of the state of alertness of the animal. We can wonder if choosing 7, 8, or 9 clusters, would have highlighted other neuron behaviors. A part of the answer is provided by figure 4, which shows that with 6 clusters, similar behaviors re-appear suggesting that a larger number of clusters would have been non-informative. As we collected multiunit recordings composed of 2-6 shapes of action potentials, it is possible that more specific behaviors might have been missed in our analyses. This is potentially the case at the cortical level where a large number of cell types have been described (Ascoli et al., 2008; DeFelipe et al., 2013) and also in the cochlear nucleus (Cant and Benson, 2003; Kuenzel, 2019). However, based on the output of small groups of neurons, five neuronal behaviors seem to be present in noise at all the levels of the auditory system.

### Noise-type sensitivity is present, in small proportions, at each stage of the auditory system but mostly in the inferior colliculus and thalamus

Ni and colleagues (2017) found about two-thirds of cortical neurons switching category from one background noise to another, suggesting that the majority of cortical neurons have a behavior specific to the type of noise. Although we initially found between 40 and 60% of such neurons in the different auditory structures, the bootstrap procedure indicated that more realistic percentages should be much lower, potentially between 10-30%. Also, the response variability, which is probably much larger in awake than in anesthetized animals, can explain the difference between our results and those of Ni and colleagues (2017). Here, these neurons were detected in auditory cortex but were found in higher proportions in subcortical structures. This indicates that only a small fraction of neurons display a behavior specific to a particular noise. We preferred to call this phenomenon noise-type sensitivity rather context-dependence (proposed by Ni and colleagues, 2017) because the latter refers to situations where the same stimulus is presented in different contexts; whereas here inserting target stimuli in two types of noise generated different auditory streams. As mentioned by Ni and colleagues (2017), if a larger number of noise types would have been tested, the proportion of neurons within each category would have been different. For example, a larger fraction of neurons can potentially be considered as target-specific or masker-specific, because masker-specific neurons in a particular type of noise can be the target-specific ones in another noise. Their assumption may still be valid but we show that it only concerns a small fraction of neurons.

### General conclusion

Over the last years, several mechanisms have been proposed to explain the robust cortical representation of speech in noise: from an ultra-sparse cortical representation (Asari et al., 2006; Schneider and Woolley, 2013), to a dynamic model of synaptic depression combined with a feedback gain normalization (Mesgarani et al., 2014), or to an adaptation to the noise statistics (Rabinowitz et al., 2013). Quantifications of high gamma activity (which reflects the average firing rate of nearby neurons, e.g. see Steinschneider et al., 2011) has recently revealed that the human auditory cortex rapidly adapts to various type of background noises (Khalighinejad et al., 2019). More precisely, speech can be reconstructed from large-scale neuronal recordings (167 electrodes) even when the background noise regularly changes, probably because neural adaptation suppresses the representation of noise features, a mechanism that seems to be independent of the attentional focus of the listeners.

Here, we propose that the noise-robustness observed in many studies at the cortical level stems, at least partially, from subcortical mechanisms (Lesica and Groethe, 2008; Rabinowitz et al., 2013) and potentially even from adaptation mechanisms already present in auditory hair cells (such as changes in gain and kinetics, see Fettiplace and Ricci, 2003). Therefore, the auditory cortex potentially inherits adaptation from earlier levels, allowing the cortical networks to focus on higher-level processing such as classifying the target stimuli into phonetic or linguistic features (Mesgarani et al., 2014), segregating the different auditory streams (Mesgarani and Chang, 2012) integrating multimodal information (Deneux et al., 2019), and retaining behaviorally important stimuli in short term (Huang et al., 2016) or long term memory (Moczulska et al., 2013; Concina et al., 2019).

## Materials and Methods

Most of the Methods are similar as in the article Souffi and colleagues (2020).

### Subjects

These experiments were performed under the national license A-91-557 (project 2014-25, authorization 05202.02) and using the procedures N° 32-2011 and 34-2012 validated by the Ethic committee N°59 (CEEA, Comité d’Ethique pour l’Expérimentation Animale) Paris Centre et Sud). All surgical procedures were performed in accordance with the guidelines established by the European Communities Council Directive (2010/63/EU Council Directive Decree).

Extracellular recordings were obtained from 47 adult pigmented guinea pigs (aged 3 to 16 months, 36 males, 11 females) at five different levels of the auditory system: the cochlear nucleus (CN), the inferior colliculus (IC), the medial geniculate body (MGB), the primary (A1) and secondary (area VRB) auditory cortex. Animals, weighting from 515 to 1100 g (mean 856 g), came from our own colony housed in a humidity (50-55%) and temperature (22-24°C)-controlled facility on a 12 h/12 h light/dark cycle (light on at 7:30 A.M.) with free access to food and water.

Two days before the experiment, the animal’s pure-tone audiogram was determined by testing auditory brainstem responses (ABR) under isoflurane anaesthesia (2.5 %) as described in Gourévitch et al. (2009). A software (RTLab, Echodia, Clermont-Ferrand, France) allowed averaging 500 responses during the presentation of nine pure-tone frequencies (between 0.5 and 32 kHz) delivered by a speaker (Knowles Electronics) placed in the animal right ear canal. The auditory threshold of each ABR was the lowest intensity where a small ABR wave can still be detected (usually wave III). For each frequency, the threshold was determined by gradually decreasing the sound intensity (from 80 dB down to −10 dB SPL). All animals used in this study had normal pure-tone audiograms (Gourévitch et al., 2009; Gourévitch and Edeline, 2011).

### Surgical procedures

All animals were anesthetized by an initial injection of urethane (1.2 g/kg, i.p.) supplemented by additional doses of urethane (0.5 g/kg, i.p.) when reflex movements were observed after pinching the hind paw (usually 2-4 times during the recording session). A single dose of atropine sulphate (0.06mg/kg, s.c.) was given to reduce bronchial secretions and a small dose of buprenorphine was administrated (0.05mg/kg, s.c.) as urethane has no antalgic properties. After placing the animal in a stereotaxic frame, a craniotomy was performed and a local anesthetic (Xylocain 2%) was liberally injected in the wound.

For auditory cortex recordings (area A1 and VRB), a craniotomy was performed above the left temporal cortex. The dura above the auditory cortex was removed under binocular control and the cerebrospinal fluid was drained through the cisterna to prevent the occurrence of oedema. For the recordings in MGB, a craniotomy was performed the most posterior part of the MGB (8mm posterior to Bregma) to reach the left auditory thalamus at a location where the MGB is mainly composed of its ventral, tonotopic, part (Redies et al., 1989, Edeline et al., 1999; Anderson et al., 2007; Wallace et al., 2007). For IC recordings, a craniotomy was performed above the IC and portions of the cortex were aspirated to expose the surface of the left IC. For CN recordings, after opening the skull above the right cerebellum, portions of the cerebellum were aspirated to expose the surface of the right CN (Paraouty et al., 2018).

After all surgery, a pedestal in dental acrylic cement was built to allow an atraumatic fixation of the animal’s head during the recording session. The stereotaxic frame supporting the animal was placed in a sound-attenuating chamber (IAC, model AC1). At the end of the recording session, a lethal dose of Exagon (pentobarbital >200 mg/kg, i.p.) was administered to the animal.

### Recording procedures

Data from multi-unit recordings were collected in 5 auditory structures, the non-primary cortical area VRB, the primary cortical area A1, the medial geniculate body (MGB), the inferior colliculus (IC) and the cochlear nucleus (CN). In a given animal, neuronal recordings were only collected in one auditory structure.

Cortical extracellular recordings were obtained from arrays of 16 tungsten electrodes (ø: 33 µm, <1 MΩ) composed of two rows of 8 electrodes separated by 1000 µm (350 µm between electrodes of the same row). A silver wire, used as ground, was inserted between the temporal bone and the dura matter on the contralateral side. The location of the primary auditory cortex was estimated based on the pattern of vasculature observed in previous studies (Wallace et al., 2000; Gaucher et al., 2013; Gaucher and Edeline, 2015). The non-primary cortical area VRB was located ventral to A1 and distinguished because of its long latencies to pure tones (Rutkowski et al., 2002; Grimsley et al., 2012). For each experiment, the position of the electrode array was set in such a way that the two rows of eight electrodes sample neurons responding from low to high frequency when progressing in the rostro-caudal direction [see examples in figure 1 of Gaucher et al. (2012) and in figure 6A of Occelli et al. (2016)]. In the MGB, IC and CN, the recordings were obtained using 16 channel multi-electrode arrays (NeuroNexus) composed of one shank (10 mm) of 16 electrodes spaced by 110 µm and with conductive site areas of 177µm^2^. The electrodes were advanced vertically (for MGB and IC) or with a 40° angle (for CN) until evoked responses to pure tones could be detected on at least 10 electrodes.

All thalamic recordings were from the ventral part of MGB (see above surgical procedures) and all displayed latencies < 9ms. At the collicular level, we distinguished the lemniscal and non-lemniscal divisions of IC based on depth and on the latencies of pure tone responses. We excluded the most superficial recordings (until a depth of 1500µm) and those exhibiting latency >= 20ms in an attempt to select recordings from the central nucleus of IC (CNIC). At the level of the cochlear nucleus, the recordings were collected from both the dorsal and ventral divisions.

The raw signal was amplified 10,000 times (TDT Medusa). It was then processed by an RX5 multi-channel data acquisition system (TDT). The signal collected from each electrode was filtered (610-10000 Hz) to extract multi-unit activity (MUA). The trigger level was set for each electrode to select the largest action potentials from the signal. On-line and off-line examination of the waveforms suggests that the MUA collected here was made of action potentials generated by a few neurons at the vicinity of the electrode. However, as we did not used tetrodes, the result of several clustering algorithms (Pouzat et al., 2002; Quiroga et al., 2004; Franke et al., 2015) based on spike waveform analyses were not reliable enough to isolate single units with good confidence. Although these are not direct proofs, the fact that the electrodes were of similar impedance (0.5-1MOhm) and that the spike amplitudes had similar values (100-300µV) for the cortical and the subcortical recordings, were two indications suggesting that the cluster recordings obtained in each structure included a similar number of neurons. Even if a similar number of neurons were recorded in the different structures, we cannot discard the possibility that the homogeneity of the multi-unit recordings differ between structures. By collecting several hundreds of recordings in each structure, these potential differences in homogeneity should be attenuated in the present study.

### Acoustic stimuli

Acoustic stimuli were generated using MatLab, transferred to a RP2.1-based sound delivery system (TDT) and sent to a Fostex speaker (FE87E). The speaker was placed at 2 cm from the guinea pig’s right ear, a distance at which the speaker produced a flat spectrum (± 3 dB) between 140 Hz and 36 kHz. Calibration of the speaker was made using noise and pure tones recorded by a Bruel and Kjaer microphone 4133 coupled to a preamplifier BandK 2169 and a digital recorder Marantz PMD671.

The Time-Frequency Response Profiles (TFRP) were determined using 129 pure-tones frequencies covering eight octaves (0.14-36 kHz) and presented at 75 dB SPL. The tones had a gamma envelop given by 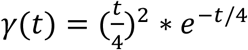,where t is time in ms. At a given level, each frequency was repeated eight times at a rate of 2.35 Hz in pseudorandom order. The duration of these tones over half-peak amplitude was 15 ms and the total duration of the tone was 50 ms, so there was no overlap between tones.

A set of four conspecific vocalizations was used to assess the neuronal responses to communication sounds. These vocalizations were recorded from animals of our colony. Pairs of animals were placed in the acoustic chamber and their vocalizations were recorded by a Bruel & Kjaer microphone 4133 coupled to a preamplifier B&K 2169 and a digital recorder Marantz PMD671. A large set of whistle calls was loaded in the Audition software (Adobe Audition 3) and four representative examples of whistle were selected. As shown in figure 1a (lower panels), despite the fact the maximal energy of the four selected whistle was in the same frequency range (typically between 4 and 26 kHz), these calls displayed slight differences in their spectrograms. In addition, their temporal (amplitude) envelopes clearly differed as shown by their waveforms (Fig. 1a, upper panels).

The four whistles were also presented in two frozen noises ranging from 10 to 24,000 Hz. To generate these noises, recordings were performed in the colony room where a large group of guinea pigs were housed (30-40; 2-4 animals/cage). Several 4-seconds of audio recordings were added up to generate the "chorus noise", which power spectrum was computed using the Fourier transform. This spectrum was then used to shape the spectrum of a white Gaussian noise. The resulting vocalization-shaped stationary noise therefore matched the "chorus-noise" audio spectrum, which explains why some frequency bands were over-represented in the vocalization-shaped stationary noise. Figures 1b et 1c display the spectrograms of the four whistles in the vocalization-shaped stationary noise (1b) and in the chorus noise (1c) with a SNR of +10 dB SPL, 0 dB SPL, −10 dB SPL. The last spectrograms of these two figures represent the noises only.

### Experimental protocol

As inserting an array of 16 electrodes in a brain structure almost systematically induces a deformation of this structure, a 30-minutes recovering time lapse was allowed for the structure to return to its initial shape, then the array was slowly lowered. Tests based on measures of time-frequency response profiles (TFRPs) were used to assess the quality of our recordings and to adjust electrodes’ depth. For auditory cortex recordings (A1 and VRB), the recording depth was 500-1000 µm, which corresponds to layer III and the upper part of layer IV according to Wallace and Palmer (2008). For thalamic recordings, the NeuroNexus probe was lowered about 7mm below pia before the first responses to pure tones were detected.

When a clear frequency tuning was obtained for at least 10 of the 16 electrodes, the stability of the tuning was assessed: we required that the recorded neurons displayed at least three successive similar TFRPs (each lasting 6 minutes) before starting the protocol. When the stability was satisfactory, the protocol was started by presenting the acoustic stimuli in the following order: We first presented the four whistles at 75 dB SPL in their natural versions (in quiet), followed by their masked versions presented against the chorus and the vocalization-shaped stationary noise at 65, 75 and 85 dB SPL. Thus, the level of the original vocalizations was kept constant (75 dB SPL), and the noise level was increased (65, 75 and 85 dB SPL). In all cases, each vocalization was repeated 20 times. Presentation of this entire stimulus set lasted 45 minutes. The protocol was re-started either after moving the electrode arrays on the cortical map or after lowering the electrode at least by 300 µm for subcortical structures.

## Data analysis

### Quantification of responses to pure tones

The TFRP were obtained by constructing post-stimulus time histograms for each frequency with 1 ms time bins. The firing rate evoked by each frequency was quantified by summing all the action potentials from the tone onset up to 100 ms after this onset. Thus, TFRP were matrices of 100 bins in abscissa (time) multiplied by 129 bins in ordinate (frequency). All TFRPs were smoothed with a uniform 5×5 bin window.

For each TFRP, the Best Frequency (BF) was defined as the frequency at which the highest firing rate was recorded. Peaks of significant response were automatically identified using the following procedure: A positive peak in the TFRP was defined as a contour of firing rate above the average level of the baseline activity plus six times the standard deviation of the baseline activity. Recordings without significant peak of responses or with inhibitory responses were excluded from the data analyses.

### Quantification of responses evoked by vocalizations

The responses to vocalizations were quantified using two parameters: (i) The firing rate of the evoked response, which corresponds to the total number of action potentials occurring during the presentation of the stimulus minus spontaneous activity; (ii) the trial-to-trial temporal reliability coefficient (CorrCoef) which quantifies the trial-to-trial reliability of the response over the 20 repetitions of the same stimulus. This index was computed for each vocalization: it corresponds to the normalized covariance between each pair of spike trains recorded at presentation of this vocalization and was calculated as follows:

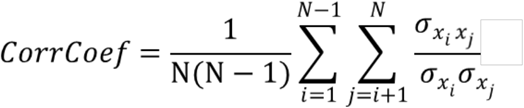

where N is the number of trials and *σx_i_x_j_* is the normalized covariance at zero lag between spike trains x_i_ and x_j_ where i and j are the trial numbers. Spike trains x_i_ and x_j_ were previously convolved with a 10-ms width Gaussian window. Based upon computer simulations, we have previously shown that this CorrCoef index is not a function of the neurons’ firing rate (Gaucher et al., 2013a).

We have computed the CorrCoef index with a Gaussian window ranging from 1 to 50 ms to determine if the selection of a particular value for the Gaussian window influences the difference in CorrCoef mean values obtained in the different auditory structures. Based upon the responses to the original vocalizations, we observed that the relative ranking between auditory structures remained unchanged whatever the size of the Gaussian window was. Therefore, we kept the value of 10 ms for the Gaussian window as it was used in several previous studies (Aushana et al., 2018; Gaucher and Edeline, 2015; Gaucher et al., 2013a; Huetz et al., 2009).

### Quantification of mutual information from the responses to vocalizations

The method developed by Schnupp and colleagues (2006) was used to quantify the amount of information (Shannon, 1948) contained in the responses to original vocalizations. This method allows quantifying how well the vocalization’s identity can be inferred from neuronal responses. Here, “neuronal responses” refer either to (i) the spike trains obtained from a small group of neurons below one electrode (for the computation of the individual Mutual Information, MI_Individual_). As this method is exhaustively described in Schnupp and colleagues (2006) and in Gaucher and colleagues (2013a), we only present below the main principles.

The method relies on a pattern-recognition algorithm that is designed to “guess which stimulus evoked a particular response pattern” (Schnupp et al., 2006) by going through the following steps: From all the responses of a cortical site to the different stimuli, a single response (test pattern) is extracted and represented as a PSTH with a given bin size (8-ms bin size was considered as in Souffi and colleagues where we showed that the optimal bin size was 8 ms for all structures). Then, a mean response pattern is computed from the remaining responses (training set) for each stimulus class. The test pattern is then assigned to the stimulus class of the closest mean response pattern. This operation is repeated for all the responses, generating a confusion matrix where each response is assigned to a given stimulus class. From this confusion matrix, the Mutual Information (MI) is given by Shannon’s formula:

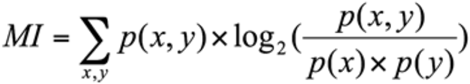

where x and y are the rows and columns of the confusion matrix, or in other words, the values taken by the random variables “presented stimulus class” and “assigned stimulus class”.

In our case, we used responses to the 4 whistles and selected the first 280 ms of these responses to work on spike trains of exactly the same duration (the shortest whistle being 280 ms long). In a scenario where the responses do not carry information, the assignments of each response to a mean response pattern is equivalent to chance level (here 0.25 because we used 4 different stimuli and each stimulus was presented the same number of times) and the MI would be close to zero. In the opposite case, when responses are very different between stimulus classes and very similar within a stimulus class, the confusion matrix would be diagonal and the mutual information would tend to log2(4) =2 bits.

The MI estimates are subject to non-negligible positive sampling biases. Therefore, as in Schnupp and colleagues (2006), we estimated the expected size of this bias by calculating MI values for “shuffled” data, in which the response patterns were randomly reassigned to stimulus classes. The shuffling was repeated 100 times, resulting in 100 MI estimates of the bias (MI_bias_). These MI_bias_ estimates are then used as estimators for the computation of the statistical significance of the MI estimate for the real (unshuffled) datasets: the real estimate is considered as significant if its value is statistically different from the distribution of MI_bias_ shuffled estimates. Significant MI estimates were computed for MI calculated from neuronal responses under one electrode.

### Quantification of the Extraction Index

To evaluate the influence of noise upon neural representation of vocalizations, we quantified the amount of vocalization encoded by neurons at a particular SNR level by calculating an extraction index (EI) adapted from a similar study in songbirds (Schneider and Woolley 2013). This metric is based on the repetition-averaged peristimulus time histogram (PSTH) of neural response with a time bin of 4 ms. Different window bins of 1, 2, 4, 8, 16, and 32 ms were also evaluated, which yielded qualitatively similar results. In this manuscript, we only report results based on 4 ms time bins. Only the PSTH during the evoked activity is taken into account in this analysis.

EI was computed as follows:

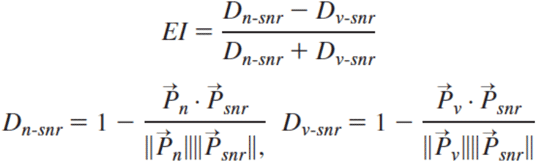

where *D_n-snr_* is the distance between PSTH 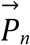 of noise and PSTH 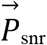 of vocalization at a NR, whereas *D_v-snr_* is the distance between PSTH 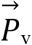 of pure vocalization and PSTH 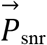 of vocalization at a particular SNR. EI is bounded between −1 and 1: a positive value indicates that the neural response to noisy vocalization is more vocalization-like, and a negative value implies that the neural response is more noise-like. The EI profile for each recording was determined by computing EI at every SNR level. The normalized inner product was used to compute distance between 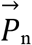 or 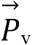 and 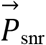, as shown in equation above.

To probe the response patterns of each neuron, we further implemented an exploratory analysis based on the calculated EI profiles as in Ni and colleagues (2017). By applying k-means clustering on the blended EI profiles from both noise conditions separately, we obtained subgroups of EI profiles, which divided the neuronal population into clusters according to the similarity of their EI profiles.

Similarity was quantified by Euclidean distance. The number of clusters was determined by the mean-squared error (MSE) of clustering, as in equation below,

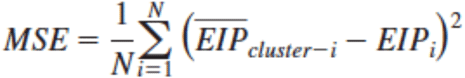

where N is the number of neurons, EIP_i_ is the EI profile of a neuron, and 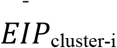 is the mean EI profile of the cluster into which this neuron is categorized.

To determine a significance level for the Extraction Index of each neuron, we generated 100 false random spike trains which follow a Poisson law based on the firing rate values obtained for each stimulus (original and noisy vocalizations). For a given SNR and recording, we computed based on these false spike trains, 100 EI_Surrogate_ values and fixed a significance level corresponding to the mean plus two standard deviations. Using this criterion, we selected only the recordings with at least one of the six EI values significantly higher than the EI_Surrogate_.

### Bootstrap procedure

To estimate the variability of the EI index generated in assigning each recording to a particular category in particular noise, we used a bootstrap strategy for all the recordings, separately for the stationary and the chorus noise. Even in anesthetized animals, auditory cortex responses can show some variability. We suspected that in a given type of noise, a recording could change category because of the response variability and/or because the border between two clusters was very close, independently to the change in noise type. From the 20 trials obtained for each stimulus during the experiment, we resampled randomly 20 trials (allowing repetitions) keeping the total number of trials the same as in the experimental data. For each resampled group of 20 trials, we recalculated both the PSTHs and the Extraction Index at each SNR then the K-means algorithm was used to define five clusters as in the experimental data. For each recording, this procedure was performed 100 times. Then, we reallocated each resampled data in the closest cluster compared to the original centroids of the experimental data to measure the percentage of changed categories relative to the original clustering.

### Statistics

To assess the significance of the multiple comparisons (masking noise conditions: three levels; auditory structure: five levels), we used an analysis of variance (ANOVA) for multiple factors to analyze the whole data set. Post-hoc pair-wise tests were performed between the original condition and the noisy conditions. They were corrected for multiple comparisons using Bonferroni corrections and were considered as significant if their p-value was below 0.05. All data are presented as mean values ± standard error (s.e.m.).

## Acknowledgments

CL and JME were supported by grants from the French Agence Nationale de la Recherche (ANR) (ANR-14-CE30-0019-01). CL was also supported by grants ANR-11-0001-02 PSL and ANR-10-LABX-0087. SS was supported by the Fondation pour la Recherche Médicale (FRM) grant number ECO20160736099 and by the Entendre Foundation. We thank Nihaad Paraouty for training us to cochlear-nucleus surgery and Jennifer Linden for insightful comments on a previous version of this article. We also wish to thank Céline Dubois, Mélanie Dumont and Aurélie Bonilla, for taking care of the guinea-pig colony.

## Competing Interests statement

The authors declare no competing financial interests.

## Notes

### Competing Interest Statement

The authors have declared no competing interest.

